# A Double Challenge: Cerebral Blood Flow During Concurrent Breath-Hold and Motor Task

**DOI:** 10.64898/2026.10.19.760082

**Authors:** Kristen Jakobsen, Martin Daiu, Hugo Critchley, Alessandro Colasanti, Iris Asllani

## Abstract

Challenging the brain with concomitant global and local demands may reveal functional constraints that are less apparent when each challenge is applied alone. We tested whether breath-holding, a global vascular challenge, modifies the local cerebral blood flow (CBF) response to motor activation.

We acquired pseudo-continuous arterial spin labelling (pCASL) MRI data at 3 T from 14 healthy volunteers during sequential finger tapping with normal breathing, finger tapping during breath-holding, and breath-holding alone. Each condition began with a resting baseline, followed by eight 25-second task blocks separated by 45-second recovery periods. We assessed CBF responses voxelwise across the whole brain and within participant-specific regions of interest in the primary motor cortex (M1) and supplementary motor area (SMA).

Despite comparable resting perfusion across conditions, the CBF increase from baseline in M1 was approximately halved when finger tapping was performed during breath-holding (26.5% versus 51.3% during finger tapping alone, p < 0.001). In the SMA, the corresponding increases were 23.2% and 41.2% (p = 0.092). On the voxelwise maps, significant CBF decreases from baseline were less spatially extensive during the combined challenge than during breath-holding alone. Global grey- and white-matter perfusion showed no statistically significant changes from baseline in any condition.

Combining global vascular and local neuronal challenges revealed CBF patterns distinct from those seen with either challenge alone. The perfusion response to added demand under stress could serve as a marker of cerebrovascular dysfunction before symptoms emerge.

## 1 INTRODUCTION

The ability of the cerebral vasculature to adjust blood flow to varying physiological demands is vital to brain function and cerebrovascular health. This adaptability encompasses responses to both local neuronal activity and systemic physiological changes. Neurovascular coupling (NVC) links neuronal activity to spatially localised changes in cerebral blood flow (CBF)^1,2^, whereas systemic factors such as changes in the arterial partial pressure of carbon dioxide (PaCO_2_) can alter vascular tone and perfusion more broadly across the brain^3,4^. How these vascular responses interact may shape the brain’s ability to meet local neuronal demands during a transient global vascular challenge or under the more persistent vascular changes associated with disease.

The cerebrovascular response to CO_2_ can be characterised experimentally using hypercapnic challenges such as breath-holding or CO_2_-enriched gas administration. Compared with CO_2_ inhalation, breath-holding offers practical advantages because it does not require specialised gas-delivery equipment and can be readily incorporated into imaging protocols. During breath-holding, continued metabolic CO_2_ production in the absence of alveolar ventilation progressively raises PaCO_2_, producing cerebral vasodilation and increased CBF^5^. This physiological response underpins the use of breath-holding as a non-invasive test of cerebral perfusion reserve. The method was introduced more than three decades ago^5^ and has since been used across several neuroimaging modalities.

Challenge-based measurements are important because baseline CBF can remain normal despite reduced cerebrovascular reserve. Hemodynamic limitations may therefore become apparent only when the vasculature is required to respond. Yet single-challenge paradigms characterise one physiological perturbation at a time and cannot establish how the circulation responds when different demands coincide.

Hypercapnia and neuronal activation impose distinct demands on the cerebral circulation. Hypercapnia produces widespread vasodilation and increased CBF, whereas neuronal activation elicits a spatially specific CBF increase through NVC^2,6^. Whether these responses combine additively, compete for available vasodilatory capacity, or otherwise modify one another remains uncertain. Experimental evidence indicates that hypercapnia can alter task-evoked hemodynamic responses, although the nature and mechanisms of this interaction remain unresolved^7,8^.

We used breath-holding as a systemic vascular challenge and sequential finger tapping to engage the motor system. We measured CBF responses to each challenge alone and to both in combination. By comparing finger tapping during normal breathing and during breath-holding, we tested whether breath-holding alters the regional perfusion response to motor activation.

To quantify these responses, we used arterial spin labelling (ASL) MRI, which uses magnetically labelled arterial blood water as an endogenous tracer to estimate tissue perfusion based on tracer kinetic theory^9^. ASL provides quantitative CBF estimates in physiological units, with millimetre-scale spatial resolution and good reproducibility across repeated measurements^10^. These features make it well suited to measuring both localised task-related perfusion responses and widespread CBF changes induced by systemic physiological challenges.

The combined challenge serves as a cerebrovascular “stress test.” The brain may respond adequately to each challenge in isolation yet show a diminished perfusion response to neuronal activation when both demands coincide. A potential application of this paradigm is to test whether the response to concurrent demands can reveal early cerebrovascular dysfunction before symptoms emerge.

## 2 METHODS

### 2.1 Participants

Following pre-enrolment screening, 14 healthy volunteers (mean age 26 ± 4.8 years, 4 females) participated in the study. Exclusion criteria included a history of neurological, psychiatric, cardiovascular, or significant systemic medical disorders; standard MRI contraindications; current smoking; hypertension; and respiratory conditions that could impair breath-hold performance (e.g., asthma).

The study was approved by the Brighton and Sussex Medical School Research Governance and Ethics Committee (RGEC) (ER/BSMSA1VA/1). All participants provided written informed consent.

### 2.2 Experimental Design

Participants completed three conditions involving breath-holding alone (BH); a motor task during normal breathing (NBM); and the same motor task during breath-holding (BHM). The motor task consisted of sequential finger-to-thumb tapping paced by a 4 Hz on-screen visual metronome.

Each condition began with a four-minute resting ASL baseline, followed by eight cycles of 25 seconds of task and 45 seconds of recovery (Figure 1). During BH, participants inhaled and held their breath, then resumed self-paced breathing during fixation. During NBM, tapping was followed by rest. During BHM, participants tapped while holding their breath. Given the potential for adaptation to repeated breath holds, all participants completed the conditions in a fixed order (BHM → BH → NBM) to standardise the sequence of breath-hold exposure across participants.

**Figure 1:**
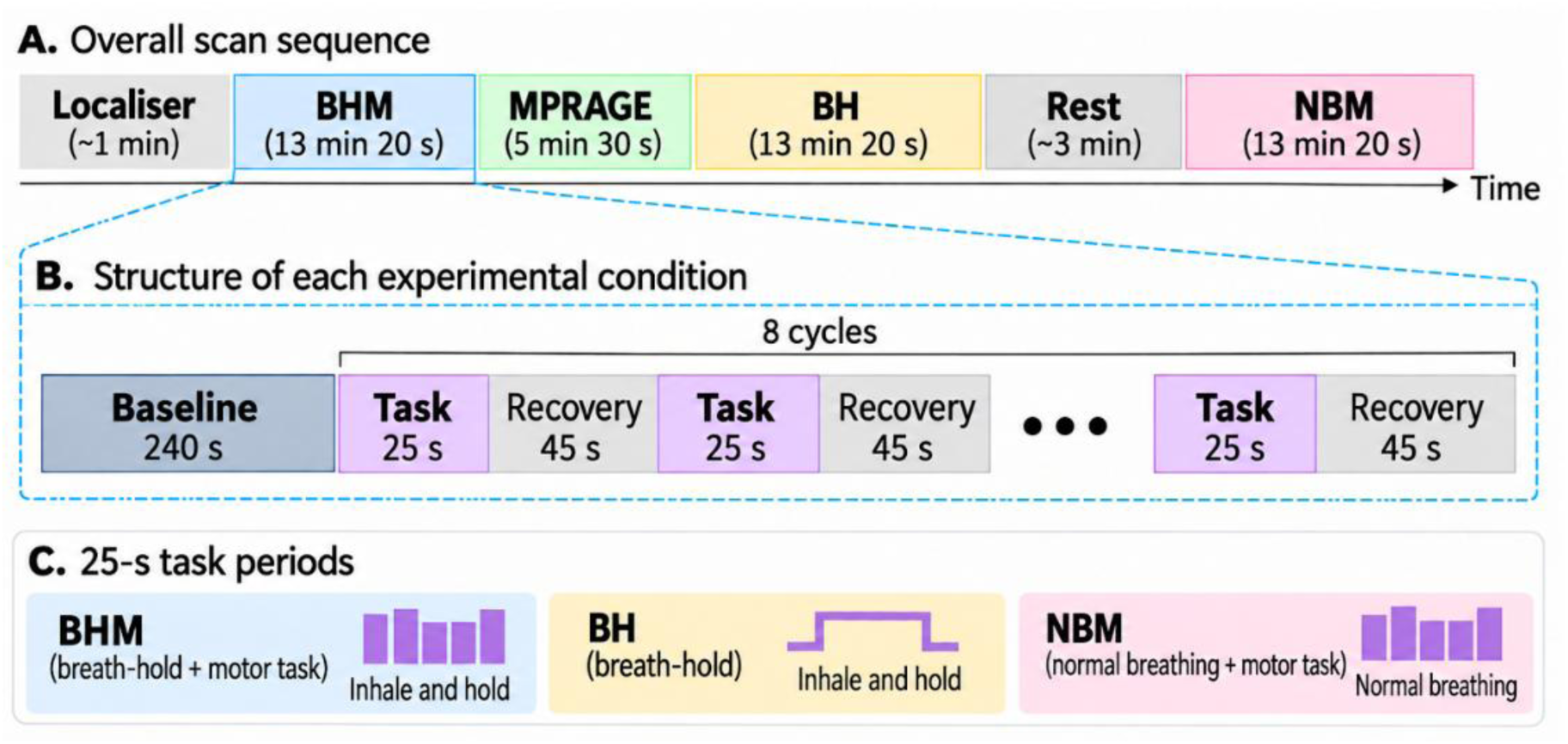
Schematic diagram of the experimental imaging protocol. Each condition consisted of a four-minute resting baseline ASL acquisition followed by the task paradigm. BHM = breath-hold + motor task; BH = breath-hold; NBM = normal breathing + motor task.

To verify that participants performed the breath-holds as instructed, the respiratory trace was acquired continuously using a BIOPAC respiration transducer (TSD221-MRI). Pulse rate was also monitored continuously using a pulse oximeter (Nonin 8600FO).

### 2.3 Data acquisition and processing

#### Imaging parameters

MRI data were acquired on a 3T Siemens PRISMA Magnetom scanner equipped with a 32-channel head coil. For each participant, an anatomical T1-weighted Magnetisation-Prepared Rapid Gradient Echo image (MPRAGE, TR = 2300 ms, TE = 2.21 ms, flip angle = 9°, FOV read = 250 mm, spatial resolution = 1-mm isotropic) was acquired for tissue segmentation and spatial normalisation purposes.

Pseudo-continuous ASL (pCASL), with background suppression^11^, was used to obtain perfusion images during baseline and tasks. 2D EPI control/label images were acquired with: TR = 5000 ms, TE = 14 ms, labelling duration = 1950 ms, post-labelling delay (PLD) = 1800 ms, slices = 20, spatial resolution = 3.4 × 3.4 × 6.0 mm. A proton-density, M0, image was also acquired using the same readout and TR as the control/label images.

#### Image pre-processing

ASL data were processed in MATLAB using custom scripts based on SPM12^12^. Motion parameters from the realignment were retained and used to generate framewise translation and rotation measures and automated outlier reports for quality control. The M0 image was resampled to the ASL image space. Pairwise perfusion-weighted difference images were computed and normalised by the M0 image to obtain relative CBF (relCBF) images. Absolute CBF was then calculated using the one-compartment model described in the ASL consensus recommendations^13^, with correction for the slice-dependent effective post-labelling delay. Individual CBF images and their mean were generated for each condition.

For spatial registration, the M0 image and associated CBF images were coregistered to the participant’s T1-weighted MPRAGE image. The MPRAGE image was segmented using SPM12, and the resulting deformation field was used to transform the ASL-derived images to MNI space at 2-mm isotropic resolution. Tissue masks were generated from the SPM tissue probability maps, retaining voxels with a combined grey matter (GM), white matter (WM), and CSF probability (c1+c2+c3) ≥ 0.30 and excluding voxels with a combined non-tissue probability (c4+c5) ≥ 0.30. The resulting masks were applied to the CBF images used for subsequent analyses.

#### Image processing

Despite motion correction, pairwise ASL difference images can contain isolated spatiotemporal outliers that may disproportionately influence CBF estimates, particularly given the intrinsically low SNR of ASL^14,15^. To reduce the influence of these outliers while preserving genuine spatial and temporal perfusion changes, we implemented a Partial Volume-Informed Robust Estimation (PVIRE) procedure (Figure 2).

**Figure 2:**
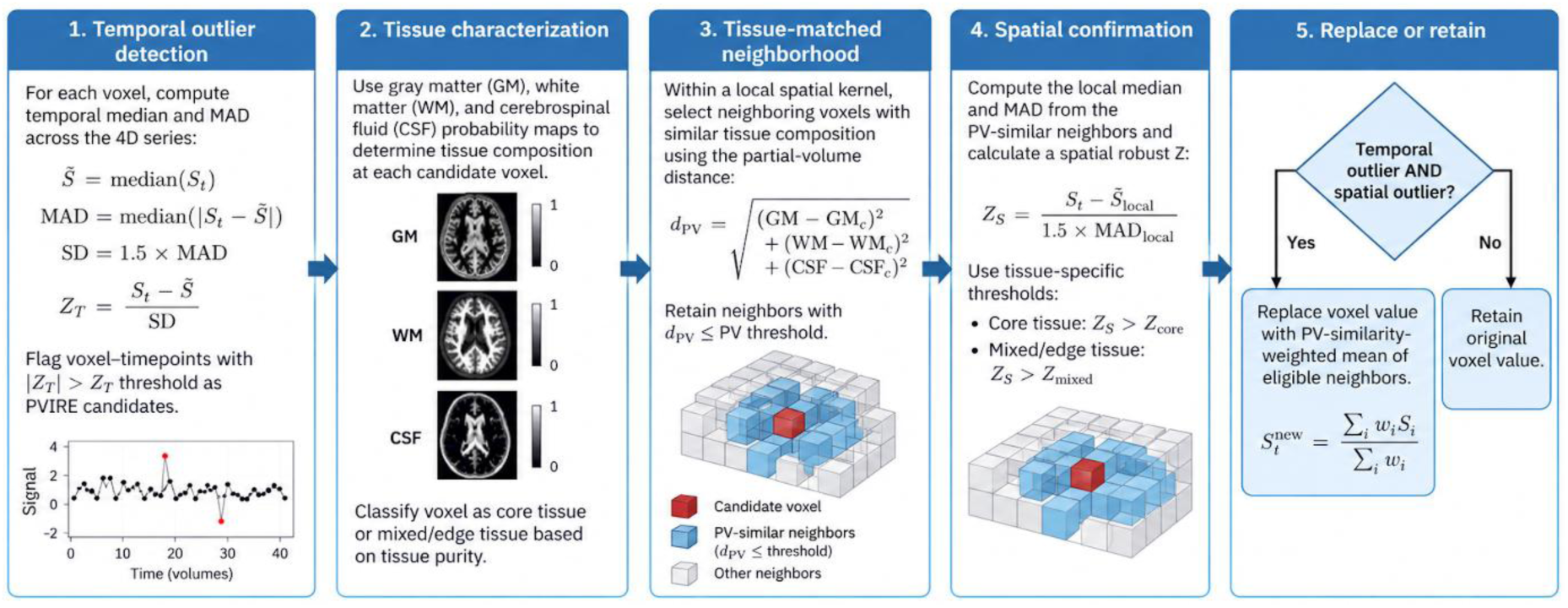
Schematic overview of the Partial Volume-Informed Robust Estimation (PVIRE) denoising procedure. For each voxel, temporal outliers are first identified from the ASL difference images using the temporal median and median absolute deviation (MAD), with the robust standard deviation estimated as 1.5 × MAD. Voxels exceeding the temporal Z-threshold are then evaluated according to their GM, WM, and CSF composition. Within a local spatial neighbourhood, voxels with similar tissue composition are selected using the PV distance d_PV_.

PVIRE was designed to identify and replace only voxels that were both temporally atypical and inconsistent with neighbouring voxels of similar tissue composition, leaving all other voxel values unchanged. Candidate temporal outliers in the relCBF images were identified voxelwise using robust temporal statistics based on the median and median absolute deviation (MAD)^16,17^. For each candidate voxel, local neighbouring voxels with similar GM, WM, and CSF partial-volume composition were identified, and the candidate was further assessed relative to this tissue-matched spatial neighbourhood^18,19^. Voxels exceeding both the temporal and tissue-specific spatial criteria were replaced by a partial-volume-similarity-weighted estimate derived from neighbouring voxels; otherwise, the original voxel value was retained.

A candidate voxel is replaced only if it also exceeds the tissue-specific spatial robust Z-threshold. Replacement values are calculated as a PV-similarity-weighted mean of neighbouring voxels; otherwise, the original voxel value is retained. Thus, PVIRE performs targeted correction of temporally and spatially confirmed outliers rather than global spatial smoothing.

### 2.4 Data analyses

#### Global analysis

The equivalence of CBF measured during the baseline (BL) acquisitions preceding the BH, BHM, and NBM conditions was assessed using a custom MATLAB script implementing the two one-sided tests (TOST) procedure^20,21^. Pairwise comparisons were performed between the BL CBF measurements for BH versus BHM, BH versus NBM, and BHM versus NBM across the 14 participants.

For each comparison, the relative difference was calculated as (*A* − *B*)/[(*A* + *B*)/2]. Equivalence bounds were defined *a priori* as ±10%. Equivalence was concluded when both one-sided tests were significant at α = 0.05, corresponding to a 90% confidence interval for the mean relative difference that lay entirely within the prespecified ±10% bounds.

Global GM and WM CBF were quantified using a custom MATLAB/SPM12 script. SPM tissue probability maps were thresholded at p ≥ 0.70, and mean CBF was calculated within each tissue mask for each condition. Baseline and task values were compared using two-sided paired t-tests, with p-values Holm-corrected across the six GM and WM comparisons.

##### Voxelwise analysis

For each participant and condition, mean baseline (BL) and task CBF images were calculated by averaging the 24 baseline and 16 task ASL measurements, respectively. Voxelwise CBF change (ΔCBF) images were calculated as the difference between mean task and mean BL CBF. For each condition, participant-level ΔCBF images were entered into a group one-sample *t*-test in SPM12 to identify regions showing significant increases or decreases in CBF relative to baseline. To compare the perfusion responses during the combined challenge and breath-holding alone, we subtracted each participant’s BH ΔCBF image from their BHM ΔCBF image and entered the resulting difference images into a group one-sample *t*-test in SPM12.

Statistical significance for the group voxelwise analyses was set at p < 0.01, with a minimum cluster extent of 50 contiguous voxels.

##### Participant-specific ROI definition

To account for inter-individual variation in the spatial location of motor activation, functional ROIs were defined separately for each participant. Within each participant, condition-specific ΔCBF measurements for NBM, BHM, and BH were analysed using voxelwise one-sample *t*-tests in SPM12 to generate participant-specific T-maps, thresholded at *p* < 0.05. The NBM T-map was used for ROI definition because NBM represented motor activation during normal breathing, without the concurrent hypercapnic challenge. Thus, the NBM activation map provided an individual functional reference for localising each participant’s motor-responsive primary motor cortex (M1) and supplementary motor area (SMA).

Subject-specific ROIs were identified from the thresholded NBM T-map using a custom MATLAB/SPM12 function. Spatially contiguous suprathreshold components were first identified and filtered by cluster size. Local T-value maxima were then identified within each retained component, with nearby maxima grouped according to spatial separation. When multiple spatially distinct peak groups occurred within the same connected component, the component was partitioned by geodesic region growing constrained to the original suprathreshold extent. Thus, all resulting ROI voxels remained within the thresholded activation map. For each participant, the M1 and SMA clusters identified from the NBM T-map were then applied to the NBM, BHM, and BH data. Mean ΔCBF and the corresponding baseline CBF were measured within each ROI for each condition, and percentage CBF change was calculated as 100 × ΔCBF/BL. Using the same NBM-defined ROI across all three conditions ensured that CBF responses were sampled from the same participant-specific motor region under normal breathing, concurrent breath-holding and motor activation, and breath-holding alone.

For visualisation, individual percentage CBF changes for NBM, BHM, and BH were plotted separately for M1 and SMA together with group means and 95% confidence intervals. The primary ROI comparison between NBM and BHM was performed using two-tailed paired *t*-tests.

## 3 RESULTS

### 3.1 Physiological data

#### Pulse rate

Paired *t*-tests showed no significant change in pulse rate from baseline in any condition. Baseline and task pulse rates were 77.6 ± 20.6 and 76.8 ± 18.1 bpm for BHM (*t*(13) = −0.68, *p* = 0.512), 71.6 ± 17.7 and 72.0 ± 16.1 bpm for BH (*t*(13) = 0.33, *p* = 0.745), and 71.9 ± 16.2 and 72.8 ± 16.9 bpm for NBM (*t*(13) = 1.96, *p* = 0.072).

#### Respiratory data

During completed breath-holds, respiratory waveforms showed an end-inspiratory peak followed by a sustained plateau, with no respiratory excursions during the prescribed hold. A two-way repeated-measures ANOVA with state (baseline, rest) and condition (BHM, BH, NBM) showed a main effect of state, *F*(1, 13) = 36.99, *p* < 0.001, η_p_^2^ = 0.74. Averaged across conditions, respiratory rate was 2.0 breaths/min lower at rest (10.9 ± 3.9 breaths/min) than at baseline (12.9 ± 4.3 breaths/min).

### 3.2 Global changes in CBF

Baseline CBF was equivalent across the three experimental conditions. Mean relative differences were 3.20% for BH versus BHM (90% CI: 0.15% to 6.24%; TOST *p* = 0.00081), −0.07% for BH versus NBM (90% CI: −2.75% to 2.61%; TOST *p* < 0.00001), and −3.27% for BHM versus NBM (90% CI: −6.48% to −0.06%; TOST *p* = 0.00130). All 90% confidence intervals fell within the prespecified ±10% equivalence bounds.

Global GM and WM CBF did not differ significantly from their corresponding baseline values during BH, BHM, or NBM after correction for multiple comparisons (all corrected *p* ≥ 0.18). In GM, CBF changed from 60.2 ± 6.7 to 58.2 ± 6.5 mL/100 g/min during BH, from 57.4 ± 6.3 to 58.6 ± 6.4 mL/100 g/min during BHM, and from 59.2 ± 7.1 to 60.7 ± 8.4 during NBM. The corresponding WM values were 37.4 ± 4.6 to 36.6 ± 4.8 mL/100 g/min, 36.4 ± 4.6 to 36.7 ± 4.2 mL/100 g/min, and 37.8 ± 3.7 to 37.9 ± 4.3 mL/100 g/min, respectively.

### 3.3 Spatial patterns of CBF responses across conditions

Figure 3 shows group-mean baseline CBF alongside maps of CBF change from baseline (ΔCBF) during NBM and BHM, expressed in mL/100 g/min. Changes were visible in the cerebellum, occipital cortex, insula, thalamus, SMA, and M1, with differences in spatial extent and magnitude between NBM and BHM.

**Figure 3:**
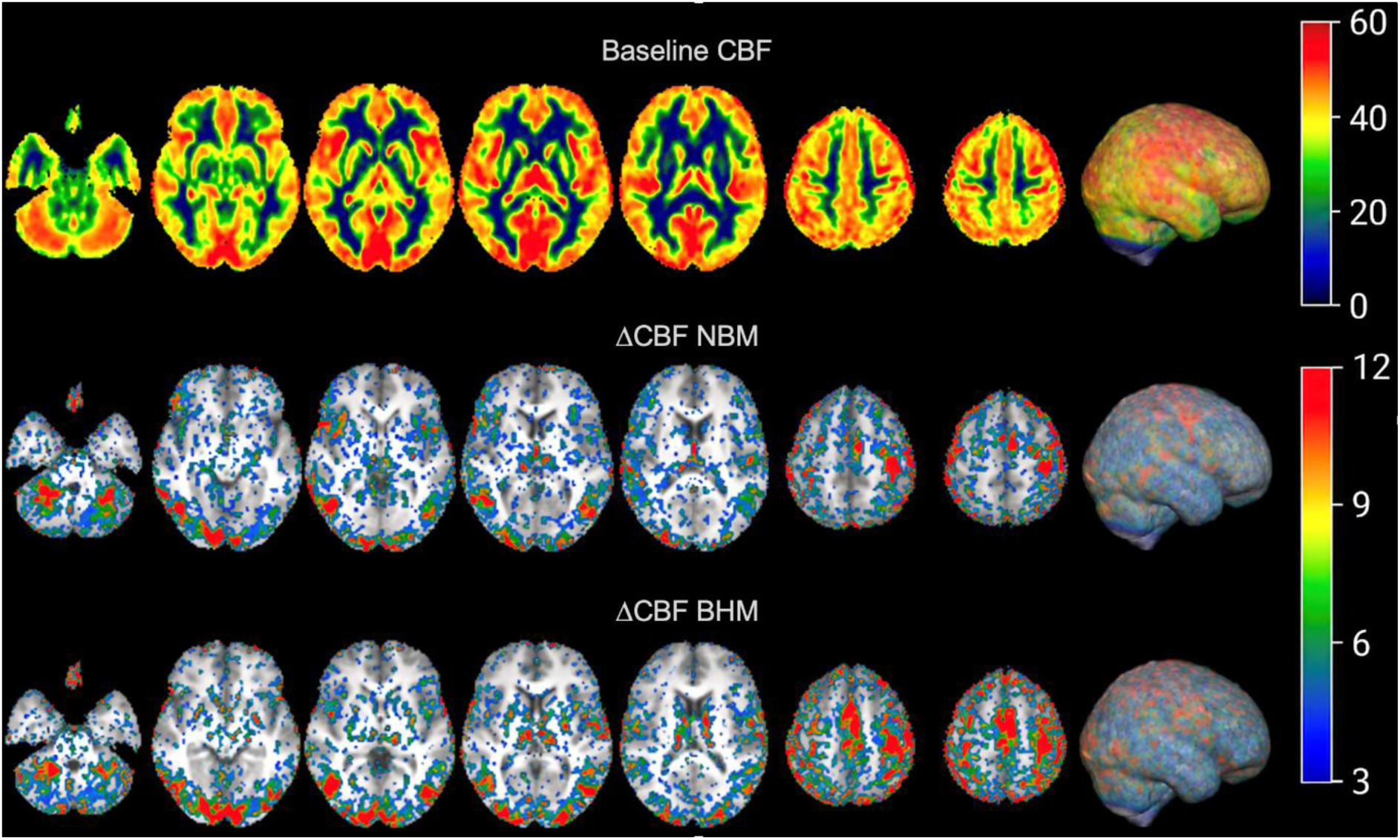
Mean ASL CBF images across participants. Baseline CBF (BL) is shown in the top row. The middle and bottom rows show condition-related changes in CBF, calculated as ΔCBF = CBFNBM − CBFBL and ΔCBF = CBFBHM − CBFBL, respectively. Colour bars indicate CBF or ΔCBF in mL/100 g/min. Axial slices are shown at MNI z-coordinates of −30, −8, 0, 6,12, 50, and 54 mm, selected to illustrate condition-related ΔCBF in the cerebellum, occipital cortex, insula, thalamus, supplementary motor area (SMA), and primary motor cortex (M1).

The thresholded *t*-maps in Figure 4 show where CBF increased during NBM and BHM (panel A) and decreased during BHM and BH (panel B), relative to each condition’s baseline. Regions with lower CBF included the thalamus and other subcortical regions, brainstem, cerebellum, medial temporal regions, and posterior cortical areas such as the occipital cortex. The reductions appeared more spatially extensive during BH than during BHM.

**Figure 4:**
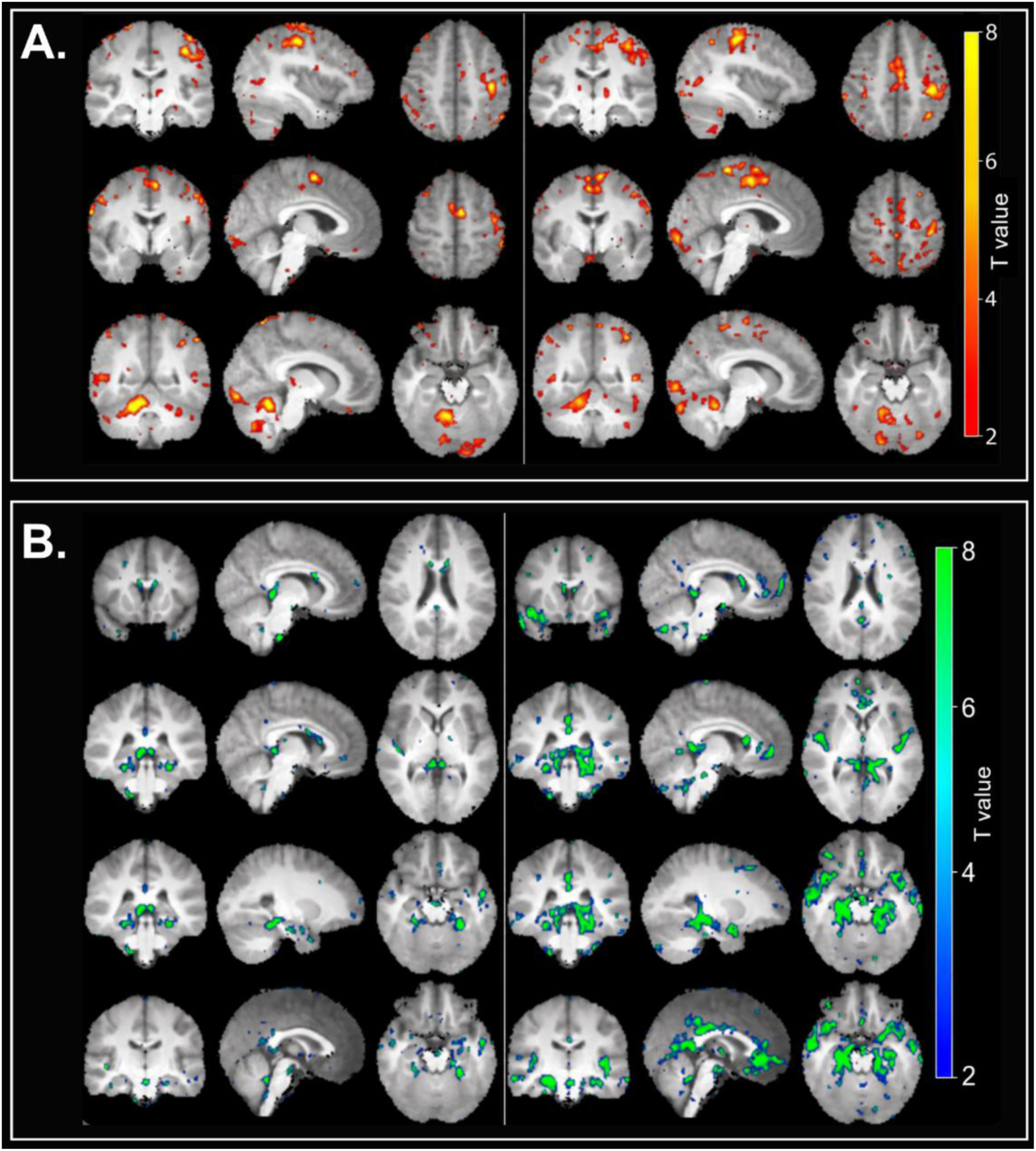
Group-level spatial patterns of condition-related CBF changes. A) T-maps for increases in CBF during NBM (left panel) and BHM (right panel) relative to baseline (BL), overlaid on the group-average T1-weighted (MPRAGE) image (*p* < 0.01, T > 2.65). Locations were selected to illustrate responses in the primary motor cortex (M1; upper row, MNI [−38, −20, 50] mm), supplementary motor area (SMA; middle row, MNI [−4, −4, 58] mm), and cerebellar and visual cortical regions (bottom row, MNI [10, −52, − 18] mm). B) T-maps showing regions in which CBF was lower during BHM (left panel) and BH (right panel) than during BL (*p* < 0.01, T > 2.65), displayed at the same locations as in panel A. No voxels survived the statistical threshold for lower CBF during NBM relative to BL.

### 3.4 Regional CBF responses

Results from the motor ROI analysis are shown in Figure 5. In M1, mean CBF change relative to baseline was 51.3% during NBM, 26.5% during BHM, and 7.2% during BH. The NBM response exceeded the BHM response by 24.8 percentage points (*t*(13) = 5.62, *p* < 0.001). In the SMA, mean CBF changes were 41.2%, 23.2%, and 5.0% during NBM, BHM, and BH, respectively. The difference between NBM and BHM did not reach statistical significance (*t*(13) = 1.82, *p*= 0.092).

**Figure 5:**
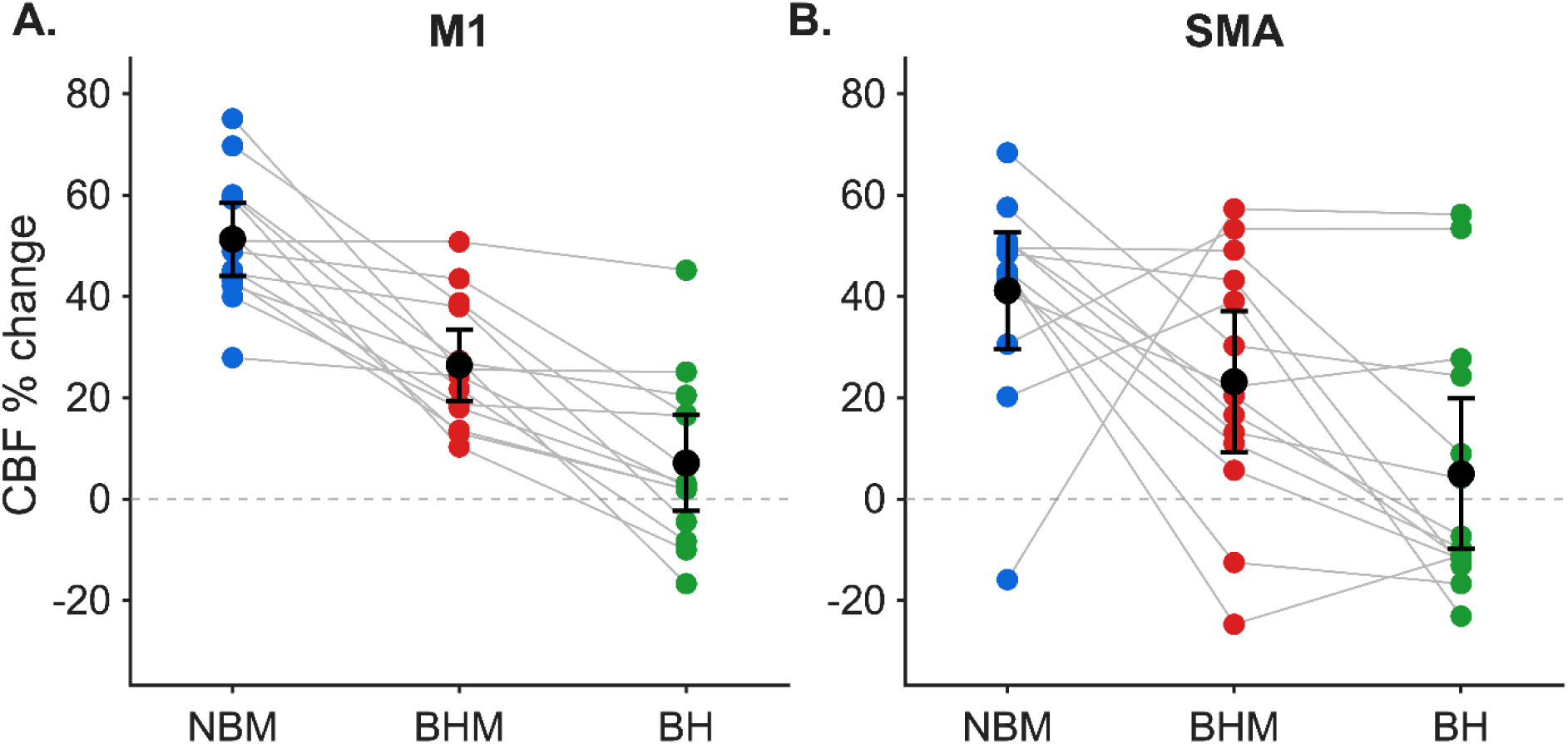
CBF responses in primary motor cortex and supplementary motor area. Percent CBF change relative to the corresponding baseline is shown for NBM, BHM, and BH in (A) primary motor cortex (M1) and (B) supplementary motor area (SMA). Participant-specific functional ROIs were defined from the NBM T-maps. Coloured points show individual participant values (NBM, blue; BHM, red; BH, green), with grey lines connecting measurements from the same participant. Black circles indicate group means and error bars represent 95% confidence intervals.

## 4 DISCUSSION

We examined whether a global vascular challenge modifies the local perfusion response to a motor task by combining breath-holding with finger tapping. The mean motor-related CBF increase in M1 fell from 51.3% during NBM to 26.5% during BHM, a significant attenuation. The estimated response was also smaller in SMA, although the difference was nonsignificant. Baseline CBF was statistically equivalent across the three conditions, and no significant changes were detected in global GM or WM CBF. The attenuation in M1 suggests a departure from simple additivity. This interpretation assumes that there were no systematic differences in the CO_2_ or blood pressure responses to breath-holding between BH and BHM.

The NBM condition established that ASL could detect robust motor-related perfusion responses using eight 25-second task blocks. CBF increased across the expected motor network, including M1, SMA, and the cerebellum. The mean M1 increase was comparable to those reported in previous block-design ASL motor studies^10,22^. Motor-related CBF increases remained detectable during BHM, showing that local functional hyperaemia can occur during a concurrent vascular challenge.

One possible explanation for the attenuated M1 response is that breath-holding shifted the vascular operating point from which the local response was generated. The rise in arterial CO_2_ expected during breath-holding promotes cerebral vasodilation^23–25^, while motor activation increases local blood flow through NVC^26,27^. If breath-holding had already recruited some of the available vasodilatory capacity, the additional perfusion response to motor activation may have been constrained. Regional differences in vascular tone, CO_2_ sensitivity, and metabolic demand could also shape the combined response^28–30^. Vasodilatory reserve was not measured directly in this study.

Earlier BOLD studies examined how vascular state influences task-evoked responses. Bruhn et al. (1994) found that visual activation remained detectable but was attenuated after acetazolamide increased baseline CBF^7^. Cohen et al. (2002) found that the local BOLD response to visual stimulation depended on global baseline CBF, complicating comparisons across regions^8^. Because BOLD reflects changes in blood flow, blood volume, and oxygen metabolism, the contribution of perfusion could not be isolated. Here, ASL provided direct estimates of CBF, with comparable baseline CBF across conditions. Using simultaneous ASL/BOLD imaging, Driver et al. (2024) detected local CBF increases in primary visual cortex during repeated breath-holding with continuous visual stimulation^31^. Their study established that local functional hyperaemia remained detectable during breath-holding and focused on validating breath-hold calibration for oxygen metabolism measurements. The comparison between NBM and BHM in the present study addresses how breath-holding changes the magnitude of the task-related perfusion response.

The coexistence of motor-related CBF increases and reductions elsewhere raises the possibility that the regional distribution of perfusion helps determine how the brain accommodates simultaneous demands. This may be particularly relevant in compromised vascular territories, where vessels can be near maximally dilated despite apparently preserved resting perfusion. During hypercapnia, further vasodilation in healthier territories can divert blood flow away from compromised regions, producing cerebral steal and potentially negative vascular reactivity^4^. A dual-challenge paradigm could complement conventional CVR measurements by assessing whether local functional demands can still be met during a concurrent vascular challenge.

The lower CBF observed in some regions during BH relative to baseline may partly reflect the timing and type of respiratory manoeuvre. Previous FAIR studies reported an immediate CBF increase during expiratory breath-holding and an initial decrease followed by an increase during inspiratory breath-holding^29,32–34^. Earlier work combining motor activation with 30-second expiratory breath-holds reported predominantly increased CBF and additive motor and vascular responses^32,33^. Our 25-second inspiratory holds may have sampled an earlier phase of the evolving response, when CBF reductions can contribute to the measured change.

Regional and interindividual differences in arterial transit time (ATT) may also have influenced the ASL measurements. Hypercapnia can shorten ATT to varying degrees across regions^23^, and the single-PLD acquisition could not separate changes in perfusion from transit-time effects. The relatively long PLD likely reduced sensitivity to modest ATT variation. ATT shortening alone is unlikely to account for the regional CBF reductions, which may reflect both the evolving physiological response and the sensitivity of ASL to changing vascular conditions.

The sample size of 14 was informed by prior ASL studies and the within-participant design. Borogovac et al. (2010) detected motor cortex perfusion increases of approximately 35% in nine participants, with mean increases of 30 and 28 mL/100 g/min across two sessions one month apart^10^. Li et al. (1999) examined motor activation, breath-holding, and their combination in six participants^35^. Subsequent studies characterised responses to 16- and 21-second breath-holds in 14 and 10 participants, respectively^25,36^. For 14 complete pairs, a two-sided paired *t*-test at α = 0.05 without multiplicity adjustment has 80% power to detect an effect size of Cohen’s *d*_z_ = 0.81 and 90% power to detect *d*_z_ = 0.94, where Cohen’s *d*_z_ is the mean within-participant difference divided by the standard deviation of those differences. The design was therefore sensitive to large within-participant effects, although power for the interaction between challenges was not established prospectively.

Conditions were administered in a fixed order to standardise prior exposure and the sequence of potential habituation across participants. With this sample size, counterbalancing would have yielded few observations per sequence for estimating order effects. Comparable baseline CBF across runs reduced concern about perfusion drift, although habituation and other order effects could not be separated from condition-specific responses. Scan times were concentrated in the early afternoon, so some time-of-day variation may also have contributed.

Even with confirmed breath-hold compliance, differences in initial lung volume, baseline CO_2_, and CO_2_ accumulation may have contributed to interindividual variability in the magnitude and timing of the CBF response. The same hold duration may therefore have sampled different phases of the physiological response across participants. Without end-tidal or arterial CO_2_ measurements, we could not separate variation in the CO_2_ stimulus from variation in cerebrovascular responsiveness. The study characterises regional perfusion responses to separate and combined challenges. Identifying the physiological mechanisms and quantifying CVR would require additional measurements.

The combined challenge revealed regional differences that were not apparent in global GM or WM CBF. Larger studies with CO_2_ monitoring and counterbalanced condition orders could clarify how individual physiology and habituation shape the combined response. Extending this paradigm to asymptomatic individuals with vascular risk factors could test whether simultaneous challenges reveal perfusion limitations while resting perfusion and responses to each challenge alone are still preserved.

## ACKNOWLEDGEMENTS

We acknowledge the support of the Clinical Imaging Sciences Centre at the University of Sussex. We thank Dr. Samira Bouyagoub for technical support, Alexandre dos Santos for assistance with data collection, and Professor Itamar Ronen for helpful feedback. We are grateful to the volunteers whose participation made this research possible.

## AUTHOR CONTRIBUTIONS

Conceptualisation: IA

Experimental design: IA, KJ, MD, AC

Data collection: KJ

Software development: IA

Image processing: IA, KJ

Formal analysis: IA

Figures: IA

Writing – original draft: IA, KJ

Writing – review and editing: All authors

Interpretation of results: All authors

## EXTERNAL FUNDING

The author(s) received no external financial support for the research, authorship, and/or publication of this article.

## AI INPUT

During manuscript preparation, IA used OpenAI ChatGPT (GPT-5.6 Sol) for code debugging, figure refinement, and language editing, and KJ used Claude (Anthropic, Sonnet 5.5) to generate initial code for processing physiological data acquired and stored in Spike2 and AcqKnowledge. All AI-assisted code was reviewed, tested, and refined by the authors, who reviewed and approved the analyses, scientific interpretations, methodological decisions, and final manuscript and take full responsibility for the work.

## ABBREVIATIONS LIST

ASL: Arterial Spin Labelling
CBF: Cerebral Blood Flow
CVR: Cerebrovascular Reactivity
NVC: Neurovascular Coupling
PaCO_2_: Arterial Carbon Dioxide Partial Pressure
BH: Breath-Hold
BHM: Breath-Hold with Motor Task
NBM: Normal Breathing with Motor Task
BL: Baseline
M1: Primary Motor Cortex
SMA: Supplementary Motor Area
PVIRE: Partial Volume-Informed Robust Estimation
relCBF: Relative Cerebral Blood Flow
ATT: Arterial Transit Time
ROI: Region of Interest
ΔCBF: Change in Cerebral Blood Flow
GM: Grey Matter
WM: White Matter
CSF: Cerebrospinal Fluid

